# A novel protein Moat prevents ectopic epithelial folding by limiting Bazooka/Par3-dependent adherens junctions

**DOI:** 10.1101/2024.03.05.583570

**Authors:** Lingkun Gu, Rolin Sauceda, Jasneet Brar, Ferdos Fessahaye, Minsang Joo, Joan Lee, Jacqueline Nguyan, Marissa Teng, Mo Weng

## Abstract

Cortical myosin contraction and cell adhesion work together to promote tissue shape changes, but how they are modulated to achieve diverse morphogenetic outcomes remains unclear. Epithelial folding occurs via apical constriction, mediated by apical accumulation of contractile myosin engaged with adherens junctions, as in Drosophila ventral furrow formation. While levels of contractile myosin correlate with apical constriction, whether levels of adherens junctions modulate apical constriction is unknown. We identified a novel Drosophila gene *moat* that maintains low levels of Bazooka/Par3-dependent adherens junctions and thereby restricts apical constriction to ventral furrow cells with high-level contractile myosin. In *moat* mutants, abnormally high levels of Bazooka/Par3-dependent adherens junctions promote ectopic apical constriction in cells with low-level contractile myosin, insufficient for apical constriction in wild type. Such ectopic apical constriction expands infolding behavior from ventral furrow to ectodermal anterior midgut, which normally forms a later circular invagination. In *moa*t mutant ventral furrow, a perturbed apical constriction gradient delays infolding. Our results indicate that levels of adherens junctions can modulate the outcome of apical constriction, providing an additional mechanism to define morphogenetic boundaries.

**Summary Statement:** Characterization of a novel gene moat demonstrates ectopic expansion of apical constriction due to abnormally high levels of Bazooka/Par3-dependent adherens junctions without defects in early patterning gene expression.

## INTRODUCTION

During morphogenesis, cell shape changes exhibit precise tissue boundaries due to tissue-specific activities of ubiquitously expressed cellular machineries, most prominently contractile myosin and cell adhesion structures (Martin and Goldstein, 2014). While contractile actomyosin generates physical force, cell adhesion often provides anchors for transmitting this force to cell cortex and maintains tissue integrity. However, how morphogenetic boundaries are defined through the activity landscape of actomyosin and cell adhesion is not well understood.

One model for such tissue-specific shape changes is the ventral furrow formation in gastrulating *Drosophila* embryos. During this process, the flat and single-layer epithelium on the ventral surface of the embryo folds inwards, serving to internalize mesoderm, the major tissue in ventral furrow. The ventral furrow region is patterned by the expression of transcription factors Snail (Sna) and Twist (Twi) (Leptin and Grunewald, 1990). Sna is a conserved transcription factor promoting epithelial-mesenchymal transition in mesoderm at a later stage (Thiery et al., 2009). However, during ventral furrow formation Sna also functions to activate contractile myosin (Manning et al., 2013).

However, not all cells in Sna-expressing ventral cells participate in ventral furrow formation. This zone is thought to cover at least three tissues from anterior to posterior (Figure 1A): the ectoderm-origin anterior midgut (we refer to it as “ectoAMG”), the endoderm-origin anterior midgut (we refer to it as “endoAMG”), and the mesoderm (Reuter and Leptin, 1994a; Technau and Campos-Ortega, 1985; Hartenstein et al., 1985). While endoAMG and mesoderm form the ventral furrow, ectoAMG is not involved. The lack of infolding in ectoAMG is due to the expression of patterning gene *huckebein* (*hkb*), a terminal gap gene that inhibits certain functions of Sna at embryo anterior (Reuter and Leptin, 1994a). However, it remains unknown how the activities of contractile myosin and cell adhesion differ between these tissues and how this leads to morphogenetic differences.

**Fig. 1.**
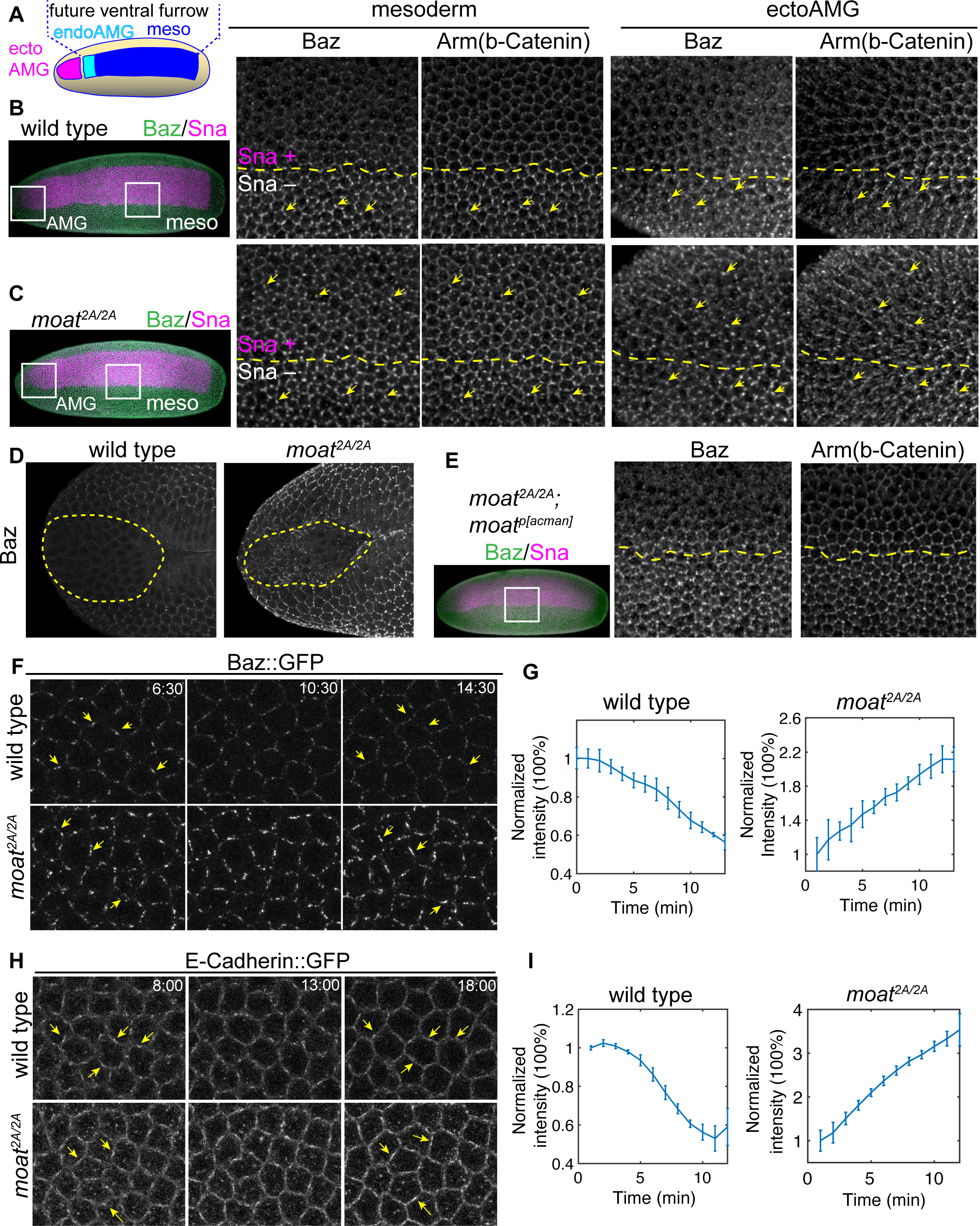
Moat disrupts the Sna-dependent downregulation of junctional Baz and adherens junctions. (A) Diagram illustrating different segments within Sna-expressing zone of early embryo. (B, C) Late stage 5, wild type (upper panels) and *moat^2A/2A^* (lower panels) embryos were immunostained for Baz, arm and Sna. Left: whole embryo’s max projection images; Right: enlarged images of boxed regions showing mesoderm/ectoAMG and their neighboring ectoderm. Yellow dashed lines: the boundaries between Sna-positive and -negative cells. Yellow arrows: Baz puncta and their corresponding junction puncta. (D) Late stage 6 wild type and *moat^2A/2A^* embryos showing junctional Baz in ectoAMG. The dashed lines: ectoAMG. (E) Genomic fragment-rescued *moat^2A/2A^* embryos. Right panel: enlarged image for the boxed region. (F) Still images from time-lapse movies of Baz::GFP in mesoderm. Arrows: the same cell edges in early (left panel) and late (right panel) time points. (G) Quantification of Baz::GFP cluster intensity from the time-lapse movies. N = 3 embryos; ∼9x15 cells per embryo. Mean and standard deviation (S.D.) are plotted. (H) Still images from time-lapse movies of E-Cad::GFP in mesoderm. Arrows: the same cell edges in early (left panel) and late (right panel) time points. (I) Quantification of E-Cadherin cluster intensity from the time-lapse movies. N = 3 embryos; ∼9x15 cells in each embryo. Mean and S.D. are plotted. For G and I, only signals from Baz or junction puncta were quantified (Fig. S1A).

The ventral furrow forms through contractile myosin-mediated apical constriction. The ubiquitously expressed non-muscle myosin II is specifically activated on apical cortices of ventral furrow cells through two parallel pathways. Components of these pathways, such as *mist*, *fog*, and *T48,* are expressed under the control of Sna and Twi (Manning et al., 2013)(Costa et al., 1994)(Kölsch et al., 2007). In contrast to tissues where myosin forms junctional cables to define tissue boundaries (Röper, 2012)(Chung et al., 2017)(Yu et al., 2021), ventral furrow cells mainly organize contractile myosin into a network on medial apical cortex which constricts apical area (Martin et al., 2009). This apical constriction induces cell and tissue shape changes, leading to the infolding of ventral furrow epithelium. The activity of myosin is patterned as a gradient along the ventral-lateral axis: high in cells around the ventral midline and low in the more flanking mesoderm (Heer et al., 2017). This gradient is important for a coordinated ventral furrow formation. The lack of infolding in ectoAMG suggests myosin activation may also be patterned along the anterior-posterior axis of the Sna-expressing zone.

The second crucial component for apical constriction is adherens junctions, the E-Cadherin(E-Cad)-catenin based cell-cell junctions that physically connect neighbor cells. Adherens junctions play multiple essential roles for morphogenetic events. One key function is to serve as anchors for contractile myosin, transmitting force to cell cortex (Martin et al., 2010)(Sawyer et al., 2009)(Roh-Johnson et al., 2012). During fly ventral furrow formation, adherens junctions connect actomyosin filaments into a tissue-wide network. Additionally, adherens junctions promote cell shape changes without myosin when differentially positioned (Wang et al., 2012). Lastly, adherens junctions are often required to maintain tissue integrity and prevent tissue rupture caused by myosin-generated tension (Martin et al., 2010)(Razzell et al., 2018).

While adherens junctions are essential for various morphogenetic events, it remains uncertain how the level of adherens junctions plays a role in modulating the boundaries of cell shape changes. Cadherin-Catenin complexes interact laterally to form clusters, which are the basic unit of adherens junctions (Strale et al., 2015)(Wu et al., 2015). Here, we define the level of adherens junctions by combining the number, size, and density of the lateral clusters of Cadherin-Catenin complexes. In early fly embryos, adherens junction clusters form many micron-sized puncta called spot adherens junctions (Tepass and Hartenstein, 1994)(Cavey et al., 2008). The number, size, and E-Cad density of these puncta change as the embryo develops. The levels of these spot adherens junctions can be patterned subcellularly to direct morphogenesis. For example, the planar polarized adherens junctions have been shown to orient the directional myosin flow and drive convergent extension (Rauzi et al., 2010).

The level of adherens junctions can also be patterned in a tissue-specific manner. Although fly ventral furrow cells require strong junctions for successful apical constriction, they are specifically patterned to start apical constriction with low levels of junctions (Weng and Wieschaus, 2017). The benefit of this patterned low level of junctions is poorly understood, but the molecular mechanism involves a Sna-dependent downregulation of junctional Bazooka (Baz), the fly polarity protein Par-3. Par-3 plays a critical role in the assembly, stability, and positioning of adherens junctions across various systems (Harris and Peifer, 2005)(Harris and Peifer, 2004)(Achilleos et al., 2010)(Simões et al., 2010). During the initial polarization of the single-layer epithelium in early fly embryo, Baz localizes to future junctional sites, which is essential for the assembly of adherens junctions (Harris and Peifer, 2005)(Harris and Peifer, 2004). However, shortly before gastrulation, junctional Baz starts to be downregulated in mesoderm, leading to low levels of adherens junctions immediately prior to apical constriction(Weng and Wieschaus, 2016)(Weng and Wieschaus, 2017). Sna is necessary and sufficient for this downregulation, but it is not clear whether the downregulation occurs in all Sna-expressing cells.

Although these mesoderm cells do regain strong junctions during apical constriction via a myosin-dependent mechanism (Weng and Wieschaus, 2016), an important question remains: does this tissue-specific reduction in junction levels immediately prior to apical constriction contribute to the spatial pattern of apical constriction and the boundaries of the ventral furrow formation? To understand the significance of Baz and junction downregulation, we undertook a genetic screen aimed at identifying mutants with aberrant Baz downregulation in ventral furrow primordium. This led to the discovery of an uncharacterized Drosophila protein, loss of which results in abnormally high levels of junctional Baz and adherens junctions. This disrupts the low level of Baz-dependent junctions patterned in Sna-expressing zone and leads to altered morphogenetic patterns inside this region.

## RESULTS

### Sna-dependent downregulation of junctional Baz is disrupted in *moat* mutants

Throughout the epithelium of fly early embryos, Baz proteins cluster into puncta at subapical cortex to assemble spot adherens junctions (Harris and Peifer, 2004). Both junctional Baz and spot adherens junctions appear as micron-size puncta and their intensity and size increase as embryos develop. However, shortly before apical constriction, junctional Baz in ventral furrow cells start to decrease in response to the rising Sna level, resulting in downregulation of spot adherens junctions (Weng and Wieschaus, 2016). This downregulation creates a pronounced contrast in the levels of junctional Baz and adherens junctions between mesoderm and lateral ectoderm (Figure 1B, mesoderm). Armadillo (Arm), the fly β-Catenin, labels spot adherens junctions as bright puncta (Fig. 1B, yellow arrow). Non-junctional Arm labels cell membrane as dim and uniform lines. Since ectoAMG also expresses Sna, we examined whether this Sna-dependent downregulation occurs in ectoAMG. Indeed, ectoAMG exhibits a similar loss of junctional Baz and adherens junctions (Figure 1B, ectoAMG). This shows that Sna’s function in downregulating junctional Baz occurs in all the Sna expressing cells. This stands in contrast to Sna’s other function: epithelial folding induced by Sna only occurs in ventral furrow but not in ectoAMG.

To understand the impact of Baz and junction downregulation on morphogenesis of this region, we identified an uncharacterized gene *CG14427* through a genetic screen looking for mutants with disrupted junctional Baz downregulation in ventral furrow. We generated a CRISPR allele of this gene deleting majority of the coding region. Homozygous mutant embryos express Sna normally but fail to downregulate junctional Baz in both mesoderm and ectoAMG, obscuring the sharp contrast in junctional Baz levels between ectoderm and these two tissues (Fig. 1C, Baz). The persistent junctional Baz leads to abnormally high levels of spot adherens junctions in both mesoderm and ectoAMG (Fig. 1C, Arm). Even at a stage when ventral furrow has fully folded and wild type ectoAMG no longer shows detectable junctional Baz, mutant ectoAMG still exhibits significant levels of junctional Baz (Fig. 1D). A small genomic fragment containing this gene restores the clear contrast in junctional Baz in mesoderm (Fig. 1E). We name this gene *moat* for its role in safeguarding the downregulation of junctional Baz in all Sna-expressing cells (*moat^2A^* is the CRISPR allele).

Live imaging shows that in wild type mesoderm, junctional Baz puncta start to diminish about 10 minutes before gastrulation (Fig. 1F, G and Movie 1) (ref). In contrast, junctional Baz puncta continues to grow in size and intensity in *moat* mutant mesoderm during the same stage (Fig. 1F, G and Movie 1). The spot adherens junctions follow a similar trend (Fig. 1H, I, Movie 2). The loss of clear contrast in junctional Baz levels between Sna-expressing cells and the lateral ectoderm could be due to elevated levels of junctional Baz in either the Sna-expressing cells only or the entire embryo. By examining junctional Baz in ectoderm, we found that *moat* mutants exhibit higher levels of junctional Baz also in the ectoderm (Fig. S1B, Movie 3). However, this global increase in junctional Baz is not due to higher expression of Baz protein. Western blot shows the amount of Baz protein in *moat* mutant and control embryos are not significantly different (Figure S1C). Thus, loss of *moat* increases the junctional localization of Baz without changing Baz protein levels.

### *moat* mutant embryos show ectopic epithelial folding of ectoAMG

Does the disruption of Baz and junction downregulation in *moat* mutant Snail-expressing zone lead to changed morphogenetic boundaries within this zone? Strikingly, we observed that the infolding behavior of ventral furrow appears to be extended to ectoAMG region in *moat* mutant embryos. Normally, the infolding is restricted to ventral furrow cells whose anterior end is at about 15 percentile egg length forming a Y-shaped structure. ectoAMG is left on embryo surface anterior to the Y-shape (Fig. 2A)(Reuter and Leptin, 1994a)(Hartenstein et al., 1985)(Technau and Campos-Ortega, 1985). However, in the *moat* mutant, the infold is extended to the very anterior end of the embryo and include the presumptive ectoAMG (Fig. 2A-B). Consistent with ectoAMG showing the most obvious defects in *moat* mutant embryo, we found that *moat* mRNA is most prominently detected in ectoAMG while its expression in other tissues is comparably more transient. Moat protein is expressed in a similar pattern and appears to associate with cell membrane (Figure S2B).

**Fig. 2.**
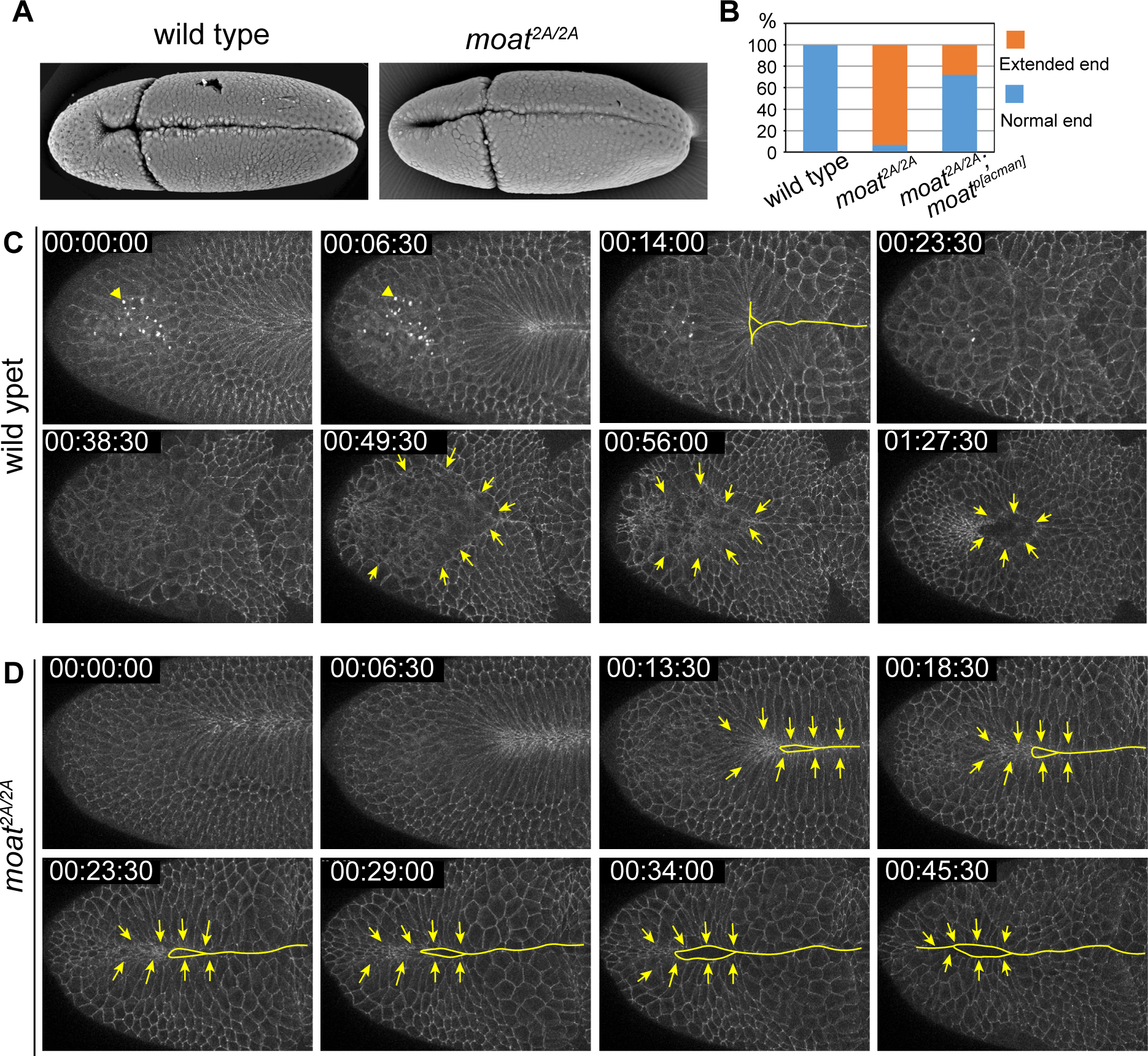
*moat* is required to prevent ectopic epithelial folding in ectoAMG. (A) SEM images of wild type and *moat^2A/2A^* embryos at stage 6. (B) Quantification of extended furrow phenotype. Wild type: N=100; *moat^2A/2A^*: N=66; *moat^2A/2A^*; Moat^p[acman]^: N=90. (C) Still images from a time-lapse movie show the internalization of wild type ectoAMG. Arrowheads in C top panels: auto-fluorescent particles from the yolk. Solid lines: indicating epithelial folds. Arrow: ectoAMG boundaries. (D) Still images from a time-lapse movie show the ectopic infolding of *moat2^A/2A^* ectoAMG. Arrows: showing directions of cell movements.

To understand the ectoAMG phenotype in *moat* mutants, we first characterized the morphogenetic course of wild type ectoAMG which has not been studied in living embryos. We used E-Cad::GFP which labels spot adherens junctions as puncta and cell membrane as dim lines. In wild type embryos, ectoAMG cells are excluded from the Y-shaped end of ventral furrow and remain on the surface of the embryo (Fig. 2C, Movie 4). About 25-30 min after the ventral furrow closure, ectoAMG starts its own course of internalization through a different mechanism. The ectoAMG boundary can be easily distinguished due to the brighter adherens junctions in neighboring ectoderm (Fig. 2C, arrows). The boundary remains mostly circular, but the encircled surface area gradually decreases. ectoAMG cells are completely internalized in about 40 min, leaving a temporary circular opening on the embryo surface (Fig. 2C, last panel). ectoAMG was thought to be internalized as part of the stomodeum invagination (Reuter and Leptin, 1994a)(Hartenstein et al., 1985)(Technau and Campos-Ortega, 1985). However, we found that they are internalized in two separate events. Stomodeum internalization is initiated later than ectoAMG. Stomodeum can be distinguished from ectoAMG by its strong adherens junctions and the formation of the characteristic triangle-shaped indentation (Movie 5). Altogether, our data indicate that the internalization of wild type ectoAMG is a morphogenetic event different from that of ventral furrow and stomodeum in both morphology and timing.

In contrast, the presumptive ectoAMG in *moat* mutants does not stay on embryo surface during ventral furrow formation but is internalized as a linear fold continuous with ventral furrow (Fig. 2D, Movie 6). Not only is this internalization earlier than that in wild type embryos, but it is also morphologically different. These mutant cells are pulled medially, perpendicular to the ventral midline, a movement similar to ventral furrow cells (Fig. 2D, arrows). This is distinct from the circular invagination of the wild type ectoAMG. In summary, *moat* mutant presumptive ectoAMG undergoes an ectopic infolding continuous with ventral furrow and as a result is internalized prematurely.

### The ectopic folding of ectoAMG in *moat* mutants is not due to aberrant expression of early patterning genes

Next, we investigated whether the ectopic infolding in *moat* mutants is due to defects in early patterning gene expression. We chose to examine two gap genes *giant* (*gt*) and *hkb* because previous studies suggest their expression patterns likely define the boundary between ectoAMG and ventral furrow cells (Eldon and Pirrotta, 1991)(Reuter and Leptin, 1994b). Since the spatial relationship between the expression of these proteins and the morphogenetic movement has not been studied at cellular levels in AMG region, we first characterized the expression of Gt and Hkb in ectoAMG and endoAMG in wild type embryos.

In wild type embryos, Gt is expressed in a narrow stripe on the ventral side of the head region, in addition to the well-studied lateral stripes (Eldon and Pirrotta, 1991). We further found that this ventral anterior Gt stripe is situated as a narrow segment in the Sna-expressing zone at blastoderm stage (Fig. S3A). To determine whether Gt is expressed in ectoAMG or endoAMG, we focused on the stage when the Y-shaped end of ventral furrow becomes apparent so we can use morphologic features to identify different tissues. Our live imaging data (Fig. 2C) show that, at this stage, wild type ectoAMG resides on embryo surface immediately anterior to the Y-shaped structure. In immunostained embryos of the same stage, we observed that the ventral anterior Gt stripe appears to surround the Y-shaped structure in max projection images, giving the impression that ectoAMG cells immediately anterior to the Y shape would be Gt(+) (Fig. 3A, upper panels, max projection). However, single optical sections reveal that the Gt(+) signals overlapping with ectoAMG in max projections are not from ectoAMG. They are from the internalized ventral furrow cells detected only in deeper optical slices (Fig. 3A, yellow arrows). ectoAMG cells, which remain on embryo surface, are all Gt(-) (Fig. 3A, encircled by yellow lines)

**Fig. 3.**
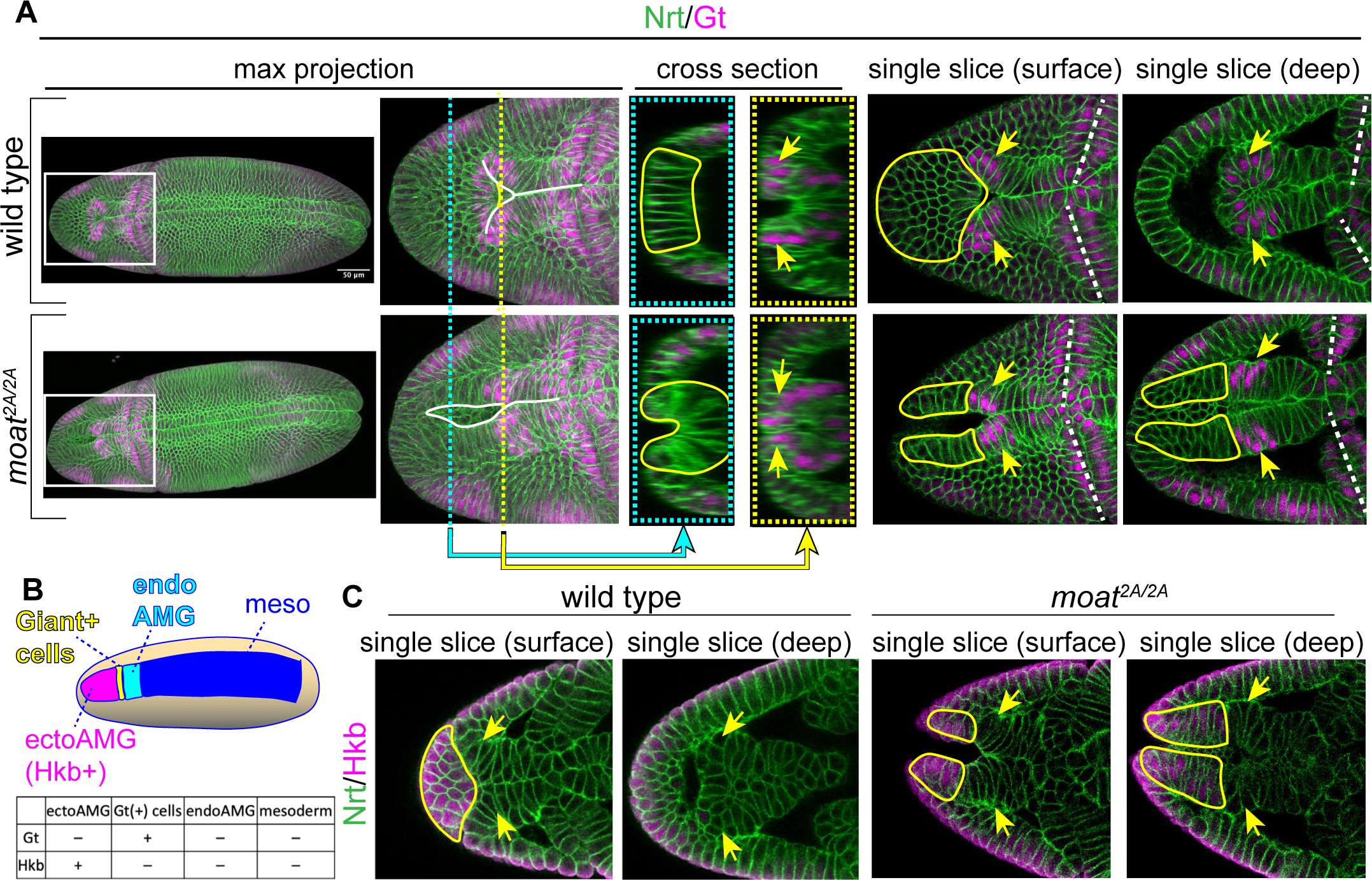
Patterning genes Gt and Hkb are expressed normally in *moat* mutant embryos. (A) wild type and *moat^2A/2A^* embryos immunostained for Gt and Nrt (membrane). White lines in max projections: anterior end of the fold. Cross-section images are transverse views (YZ) generated from confocal stacks (XYZ) using the Reslice function in Fiji at the X positions indicated by the dashed cyan and yellow lines. Single slice images are single optical sections from confocal stacks. Enclosed yellow lines: presumptive ectoAMG based on morphology and position. Yellow arrows: Gt(+) cells. White dashed lines: cephalic furrows. (B) Diagram showing four distinct tissues in Sna expression zone. (C) Single optical sections of anterior regions of embryos immunostained for Hkb and Nrt. Enclosed yellow lines: Hkb(+) ectoAMG. Arrows: presumptive Gt(+) cells.

These data indicate that Gt(+) cells are not ectoAMG but constitute the anterior end of the internalized ventral furrow. The anterior end of the ventral furrow was thought to be endoAMG based on horseradish peroxidase labeling experiments (Hartenstein et al., 1985) (Reuter and Leptin, 1994a), which would suggest the Gt(+) cells are endoAMG. However, by following Gt(+) cells in later stage embryos, we found that they do not contribute to AMG structure: they remain in the head region, and spread to surround the invaginated stomodeum (Fig S3B). This is consistent with the previous study on Gt expression (Eldon and Pirrotta, 1991). Taken together, we concluded that the anterior ventral Gt(+) cells are an uncharacterized tissue positioned as a segment in Sna-expressing zone at blastoderm stage between ectoAMG and endoAMG primordia (Fig. 3B).

Next, we examined the expression of another gap gene *hkb* in wild type embryos. We found that Hkb protein is expressed in the presumptive ectoAMG cells but not the Gt(+) cells (Fig. 3C). Single optical slices show that Hkb is detected in ectoAMG cells occupying the surface position immediately anterior to the Y-shape structure (Fig. 3C, wild type, encircled by the yellow line). However, Hkb is absent in the presumptive Gt(+) cells identified as the anterior end of internalized ventral furrow (Fig. 3C, wild type, yellow arrows). Taken together, our data showed that there are four different segments in wild type Sna expressing zone: 1) Gt(-) Hkb(+) ectoAMG; 2) the uncharacterized Gt(+) Hkb(-) cells; 3) Gt(-) Hkb(-) endoAMG; 4) Gt(-) Hkb(-) mesoderm (Fig. 3B).

In *moat* mutant embryos, neither Gt or Hkb expression pattern appears to be altered. The ventral Gt stripe remains at about 15 percentile egg length similar to that in wild type embryos. The elongated mutant furrow passes through the Gt stripe to include the anterior Gt(-) cells (Fig. 3A, *moat^2A/2A^*, max projection). These Gt(-) cells often form an incomplete tube with a narrow opening on the embryo surface which is continuous with the epithelial tube formed by ventral furrow (Fig. 3A, *moat^2A/2A^*, cyan cross section). As a result, Gt(+) cells appear as a segment of the continuous tube (Fig. 3A, *moat^2A/2A^*, deep single slice, cross sections). Similarly, *moat* mutant embryos express Hkb normally. Hkb remains to be expressed in the anterior ventral cells despite their ectopic infold in *moat* mutants (Fig. 3C, *moat^2A/2A^*, encircled by yellow lines). Hkb is also not detected in the presumptive Gt(+) cells in *moat* mutant embryos identified by their positions, same as the wild type (Fig. 3C, *moat^2A/2A^*, yellow arrows).

These data indicate that the ectopic infolding observed in *moat* mutant embryos is not due to an expansion of ventral furrow fate to the anterior end of the embryo, and ectoAMG are most likely specified properly in *moat* mutant embryos.

### ectoAMG in *moat* mutant embryos exhibit ventral furrow-like apical constriction

Because ventral furrow formation is driven by myosin-mediated apical constriction, we investigated whether the ectopic infolding of ectoAMG in *moat* mutant embryos is driven by the same mechanism. During ventral furrow formation, myosin is activated by two parallel pathways that recruits RhoGEF2 to apical cortex: one mediated by a transmembrane Gα protein Cta (Costa et al., 1994)(Barrett et al., 1997)(Dawes-Hoang, 2005)(Manning et al., 2013) and the other by an apical membrane-localized protein T48 (Kölsch et al., 2007). Eliminating both *cta* and *T48* completely abolishes apical constriction in ventral furrow.

First, we tested whether the loss of *cta* and *T48* would suppress the ectopic apical constriction of *moat* mutant ectoAMG. Indeed, in *moat, cta*, *and T48* triple mutant embryos, not only ventral furrow but also ectoAMG fail to undergo infolding and are left on the embryo surface (Fig 4A). This suggests that the ectopic infolding in *moat* mutant ectoAMG requires the same pathways that activate myosin in wild type ventral furrow.

**Fig. 4.**
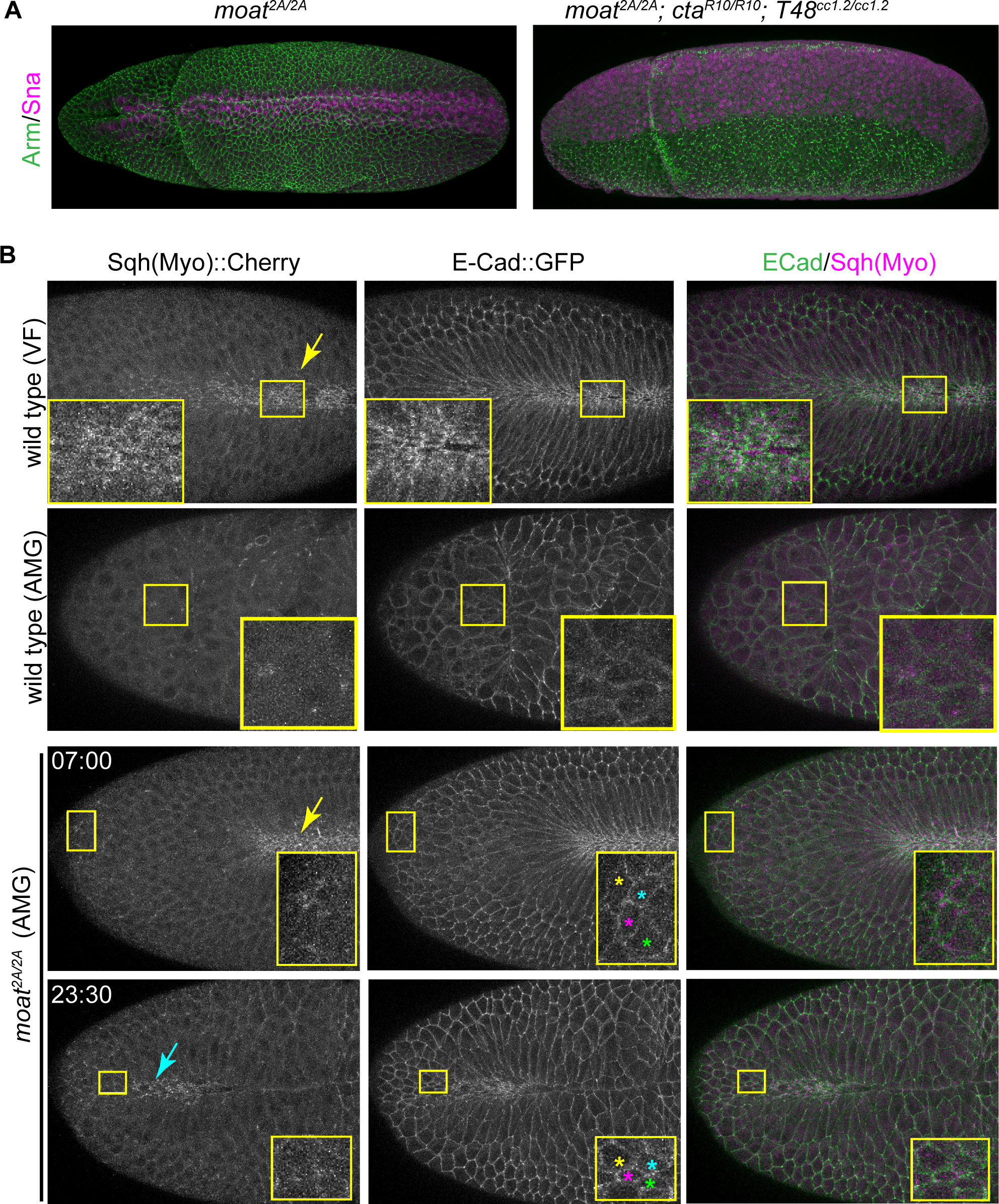
Ectopic apical constriction in *moat* mutant embryo extoAMG. (A) Immunostained embryos of stage 7. (B) Still images from time-lapse movies og indicated genotypes. Inserts: enlarged images of boxed regions. Colored stars: the tracked cells at the two time points. Yellow arrows: active myosin in either wild type or *moat^2A/2A^* ventral furrows. Cyan arrow: active myosin in highly constricted cells in *moat^2A/2A^* ectoAMG.

Next, we examined the level of contractile myosin in ectoAMG. Past studies show that contractile myosin can be detected as intense cortical myosin structures that are distinguishable from the uniformly distributed inactive myosin in the cytoplasm, and its intensity correlates with apical constriction (Xie and Martin, 2015) (Vasquez et al., 2016)(Heer et al., 2017). We compared contractile myosin in three conditions: (1) wild type ventral furrow region, known to undergo robust myosin-mediated infolding; (2) wild type ectoAMG, a tissue that does not undergo infolding; (3) *moat* mutant ectoAMG that undergoes ectopic infolding. Wild type ventral furrow cells display strong contractile myosin on the apical cortex, and undergo apical constriction and infolding (Fig. 4B, top row; Movie 7). By contrast, wild type ectoAMG cells display detectable but low levels of contractile myosin and exhibit little apical constriction (Fig. 4B, second row; Movie 8). Compared to wild type ectoAMG, *moat* mutant ectoAMG cells appear to exhibit slightly higher levels of contractile myosin, which constrict their apical surfaces: the apical areas of the tracked cells reduce as they form the ectopic fold (Fig. 4B, bottom two rows). The increase in contractile myosin is more noticeable in highly constricted cells (Fig. 4B, bottom row, cyan arrow; Movie 9). Nevertheless, the contractile myosin in *moat* mutant ectoAMG is notably weaker than that in ventral furrows of either wild type or *moat* mutant embryos (Fig. 4B, yellow arrows).

### The increase in contractile myosin in *moat* mutant ectoAMG is mild

The above comparison of contractile myosin could be confounded by apical constriction occurring only in *moat* mutant but not wild type ectoAMG cells, since contractile myosin can appear more concentrated in cells with reduced apical area. To compare myosin activities without this complication, we disrupted apical constriction by depleting myosin’s anchors to the cell cortex: adherens junctions. Without adherens junctions, apical constriction cannot occur, but myosin activity can be assessed by observing its impact on apical membrane deformation through scanning electron microscopy (SEM) (Martin et al., 2010) (Fig. 5A).

**Fig. 5.**
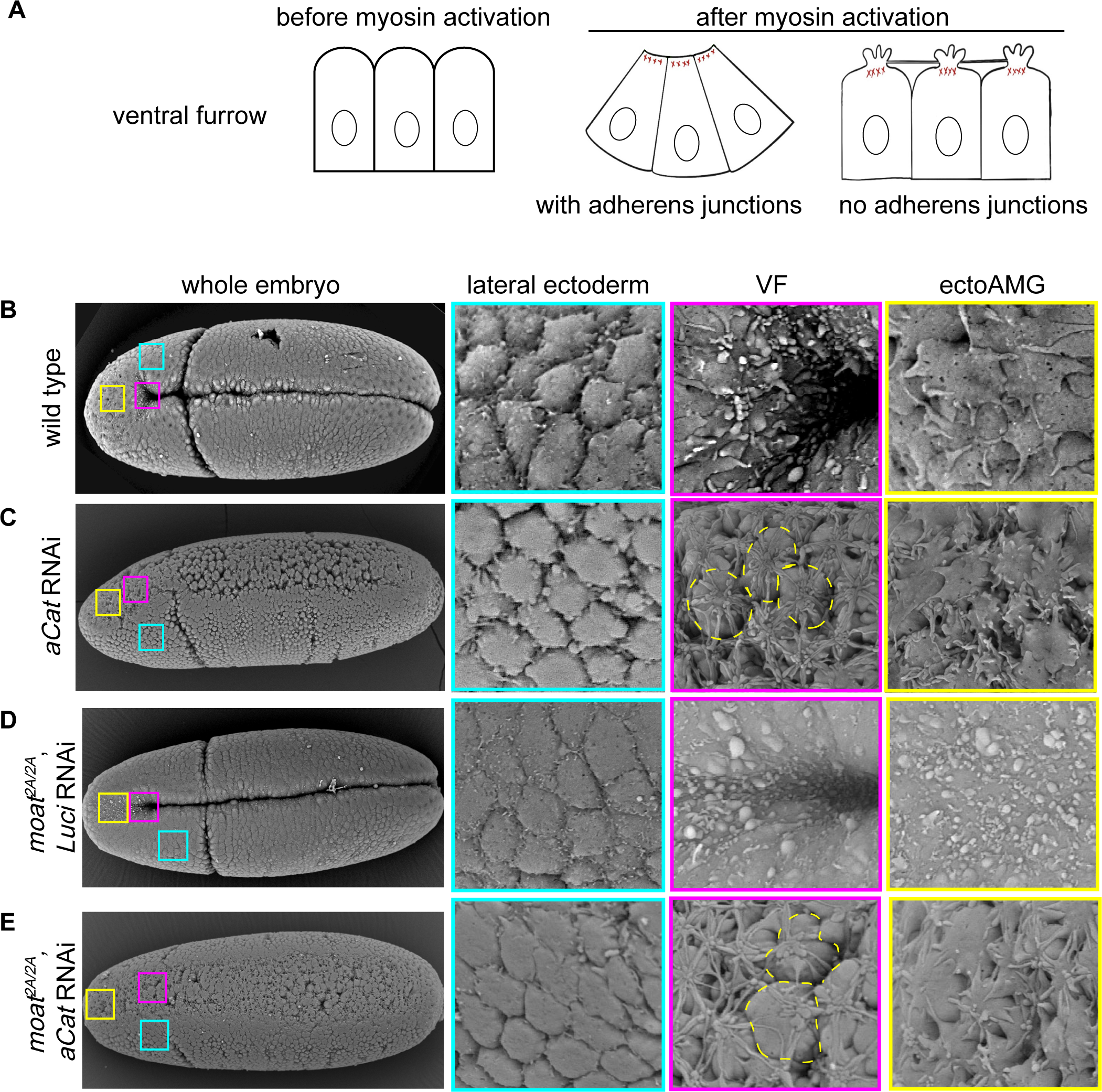
Contractile myosin in *moat* mutant ectoAMG is much weaker than that in ventral furrow. (A) Diagram illustrating the morphological changes of apical surface in response to contractile myosin. (B-E) SEM images of embryos of indicated genotypes. High magnification images of specific tissues boxed in the whole embryo displayed on the right. Yellow circles of dashed lines: individual cells ruptured from neighboring cells, displaying membrane foci and tethers.

Apical membrane deformations in response to different levels of contractile myosin have been well characterized in ventral furrow cells in both wild type and adherens junction-defective embryos (Sweeton et al., 1991) (Martin et al., 2010). Before myosin activation, all cells display smooth and dome-shaped apices with well-defined cell boundaries (Fig. S4A). As embryos develop to gastrulation, lateral ectoderm retains the dome shaped apices since myosin is not activated in this tissue (Fig. S4A-C, Fig. 5B, lateral ectoderm). However, in ventral furrow cells, apical membrane morphology shows distinct changes in response to changing levels of contractile myosin. Initially, low levels of myosin contraction lead to flattened apical surfaces resulting in less distinct cell boundaries (Fig. S4B). Later, strong myosin contraction leads to invisible cell boundaries, and accumulation of small membrane blebs (Fig. S4C, Fig. 5A, B).

In junction-defective embryos such as α-Catenin (α-Cat) RNAi embryos, the non-constricting ectoderm cells still maintain dome-shaped apices (Fig. 5C, lateral ectoderm). However, ventral furrow cells in the same embryo display a dramatic deviation from their wild type counterparts: complete ruptures occur between cell apices due to high levels of myosin contraction without adherens junctions (Fig. 5C, VF). As a result, the apical cortices in individual cells contract independently, leading to apical membrane accumulation in one focus on top of each cell. These membrane foci are connected by long membrane tethers due to the residual cell adhesion between cells (Fig. 5A, C, VF).

With these known responses of apical membrane to contractile myosin, we examined the apical membrane morphology in ectoAMG of different genotypes. First, we examined ectoAMG in wild type background with or without α-Cat RNAi. Although wild type ectoAMG cells do not undergo apical constriction, their apices appear to be flattened (Fig. 5B, ectoAMG), resembling the response of ventral furrow cells to low-level myosin contraction (Fig. S4B). This is consistent with the low-level contractile myosin in wild type ectoAMG (Fig. 4B). In α-Cat RNAi embryos, ectoAMG displays small cell tears and some membrane tethers reaching to neighbor cells, but the degree of deformation is much less than that in the ventral furrow: no membrane foci can be observed (Fig. 5C, ectoAMG vs VF). This suggests wild type ectoAMG cells have low levels of contractile myosin that leads to small cell tears when adherens junctions are depleted.

Next, we examined *moat* mutant embryos with or without α-Cat RNAi. Loss of *moat* alone does not change lateral ectoderm and ventral furrow apical membrane morphology significantly. In *moat* mutants, lateral ectoderm still shows smooth apices with well-defined cell boundaries and ventral furrow cells still show small membrane blebs with invisible cell boundaries (Fig. 5D vs 5B, lateral ectoderm, VF). Similarly, these two tissues with α-Cat RNAi in *moat* mutant background resemble their counterpart tissues with α-Cat RNAi in wild type background: ectoderm maintains smooth apices and clear cell boundaries while ventral furrow displays complete cell ruptures with membrane foci and long membrane tethers (Fig. 5E vs 5C, lateral ectoderm, VF).

By contrast, *moat* mutant ectoAMG differs significantly from its wild type counterpart in apical membrane morphology. *moat* mutant ectoAMG displays apical membrane features similar to that of ventral furrow with strong contractile myosin: unrecognizable boundaries and small membrane blebs on apical surfaces (Fig. 5D, ectoAMG; Fig. S4C). This is consistent with *moat* mutant ectoAMG exhibiting ventral furrow-like apical constriction behavior. However, *moat* mutant ectoAMG responds differently from ventral furrow when α-Cat is further removed. ectoAMG with α-Cat RNAi in *moat* mutant background lacks the complete cell rupture phenomenon observed in ventral furrow with α-Cat RNAi: there are no membrane foci on top of individual cells but only small cell tears and tethers (Fig. 5E ectoAMG vs 5C VF or 5E VF). This phenotype is more similar to ectoAMG with α-Cat RNAi in wild type background, except one detail: the small membrane tethers in *moat* mutant background appear to be straighter and reaching to neighboring cells more, suggesting moderately increased contractile myosin in *moat* mutant ectoAMG (Fig. 5E vs 5B, ectoAMG).

Taken together, we conclude that, despite the similar apical constriction behavior, the level of contractile myosin in *moat* mutant ectoAMG is much lower than that in ventral furrow. Contractile myosin in *moat* mutant ectoAMG is only moderately increased compared to wild type ectoAMG. We also showed that contractile myosin requires adherens junctions for *moat* mutant ectoAMG to exhibit the ventral furrow-like morphology such as reduction in apical area and apical membrane blebs.

### Junctional Baz is essential for ectopic apical constriction in *moat* mutant ectoAMG

How does the low level of contractile myosin in *moat* mutant ectoAMG achieve morphogenetic outcomes similar to the strong contractile myosin in ventral furrow? Although *moat* mutant ectoAMG shows less contractile myosin compared to wild type ventral furrow, it displays higher levels of junctional Baz and Baz-dependent adherens junctions (Fig. 1B,D). Since adherens junctions are required to engage contractile myosin for force transmission, we reasoned that the high levels of Baz-dependent adherens junctions may compensate for the low levels of myosin. As discussed earlier, adherens junctions in early fly embryos are formed through two mechanisms. The Baz-dependent mechanism determines the junction levels before entering apical constriction, while the myosin-dependent mechanism is essential for regaining adherens junctions in normal ventral furrow during apical constriction (Weng and Wieschaus, 2016). We hypothesized that ventral furrow formation may predominantly rely on myosin-dependent adherens junctions, while the ectopic infolding of *moat* mutant ectoAMG would require Baz-dependent adherens junctions.

Consistent with our hypothesis, we found that knocking down Baz abolishes the ectopic infolding of ectoAMG in *moat* mutant embryos and restores the anterior furrow end to the Gt(+) cells (Fig. 6A; Fig. 3A). By contrast, Baz depletion has a much milder effect on ventral furrow. Although Baz is undetectable using immunostaining, ventral furrow still forms (Fig. 6A, arrows). These phenotypes can be clearly seen using SEM. In *moat* mutant embryos with Baz RNAi, there is a clear distinction between ectoAMG and ventral furrow: ectoAMG cells are relaxed with little signs of constriction, while ventral furrow cells are strongly constricted, displaying extensive membrane blebbing (Fig 6B, arrows). The ectopic infolding of ectoAMG is lost in all *moat* mutant with Baz RNAi embryos we examined (Fig. 6C). Ventral furrows, on the other hand, largely form with Baz depleted (Fig. 6B, bottom two rows). The closure of the two ends of the ventral furrow tends to be incomplete, but more than 90% of embryos close at least the middle third of the ventral furrow (Fig. 6D).

**Fig. 6.**
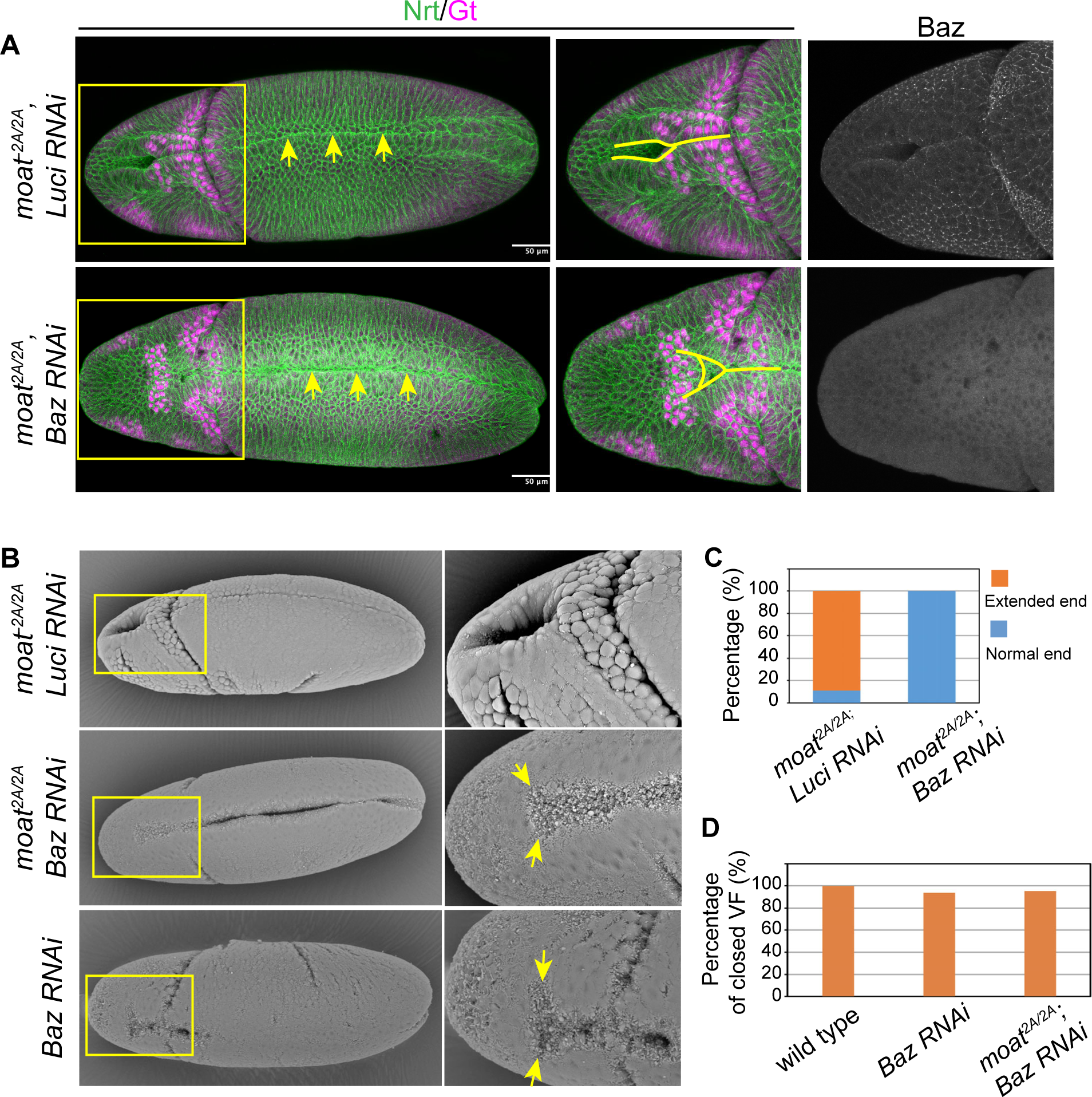
Baz is required for the ectopic infolding in *moat* mutant ectoAMG. (A) Immunostained embryos with enlarged images of the boxed regions. Arrows: closed ventral furrow; solid lines: anterior end of the fold. (B) SEM images of embryos of indicated genotypes. The enlarged images of AMG regions show the ends of the epithelial folds. (C) Quantification of normal and extended furrow end phenotypes. *moat^2A/2A^*; *luci* RNAi: N=30; *moat^2A/2A^*; *Baz* RNAi: N=20. (D) Percentage of embryos with at least middle one third of their ventral furrow closed. Wild type: N=36; *Baz* RNAi: N=32; *moat^2A/2A^*; *Baz* RNAi: N=42.

These results suggest that in ventral furrow, strong myosin contraction is mostly sufficient for generating both tension force and strong adherens junctions to mediate effective apical constriction and epithelial folding. Whereas, in *moat* mutant ectoAMG, although myosin contraction is slightly higher than that in wild type ectoAMG, it is insufficient to induce ventral-furrow-like apical constriction without the high levels of Baz-dependent adherens junctions. The combination of weak contractile myosin and strong Baz-dependent adherens junctions leads to the expansion of infolding behavior from ventral furrow to ectoAMG.

### Flanking mesoderm cells in *moat* mutant embryos undergo ectopic apical constriction

If the elevated Baz-dependent adherens junctions in *moat* mutants can enhance apical constriction in *moat* mutant ectoAMG, we asked whether other tissues with low-level contractile myosin would also undergo ectopic apical constriction in *moat* mutant embryos. We chose to examine the flanking mesoderm because of its low levels of contractile myosin and lack of apical constriction in wild type embryos. Not all mesoderm cells constrict equally. Along the ventral-lateral axis of mesoderm, there is a gradient of apical constriction resulted from a gradient of contractile myosin: cells close to the midline show stronger contractile myosin and constrict their apices, while cells further from the midline show lower contractile myosin and expand their apices (Heer et al., 2017). This gradient of constriction is essential for coordinated ventral furrow formation.

To test whether loss of *moat* leads to ectopic apical constriction in the flanking mesoderm, we tracked rows of cells from the ventral midline to quantify changes in their apical areas (Fig 7A). E-Cad::GFP was used to label adherens junctions and the cell outline for tracking individual cells. In wild type embryos, we observed a gradient of adherens junctions, stronger in cells closer to ventral midline (Fig 7B, left). This suggests a gradient of junctions is formed in response to the gradient of contractile myosin, consistent with the essential role of myosin-dependent junction mechanism (Weng and Wieschaus, 2016). This shows the wild type flanking mesoderm cells resemble ectoAMG in both low contractile myosin and low adherens junctions. By contrast, in *moat* mutants, the flanking mesoderm show significantly higher levels of adherens junctions due to high levels of junctional Baz, and the gradient of adherens junction levels becomes less pronounced (Fig 7B, right).

**Fig. 7.**
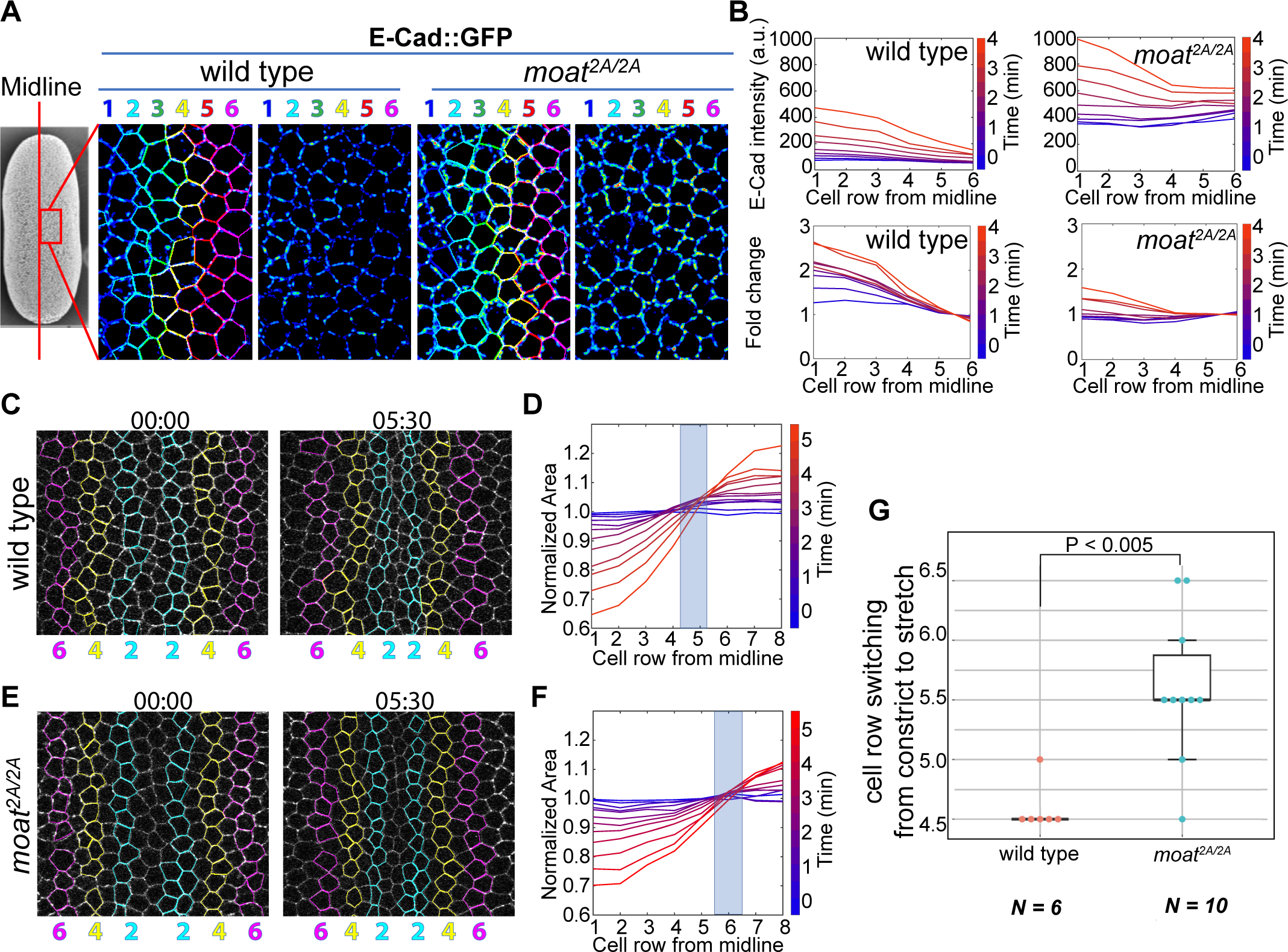
Ectopic apical constriction in flanking mesoderm cells. (A) Individual cells are tracked and assigned row numbers based on their positions relative to the ventral midline. Heatmap images show relative E-Cad intensity. (B) Top panels: the intensity of spot adherens junctions of different cell rows over time. Lower panel: the relative intensity of spot adherens junctions normalized by row 5 and 6 over time. Each curve represents junction intensities across the rows from the middle to the side. The blue-to-red color spectrum represents time points. (C, E) Still images from time-lapse movies with tracked rows of cells on both sides of the midline. Numbers and colors are consistent with those in (A). (D, F) Normalized apical area plotted against cell row number for the time-lapse movies in (C, E). Curves for individual time points are represented using the blue-to-red color spectrum. Each plot in (B, D and F) shows measurements from individual embryos, and the number of cells in each row and at each time point may vary ranging from 23 to 32. (G) Quantification of the cell row number at which cells switch from constriction to expansion. Wild type: N=6; *moat^2A/2A^*: N=10. p=0.0044 determined using student T-tests (two tails, two-sample unequal variance).

Wild type mesoderm displays a highly reproducible gradient of apical constriction (Fig. 7C, D, Movie 10). Cells in the first four rows from the midline show apical constriction while cells from row five start to show apical expansion (quantified in Fig. 7D), in agreement with the previous report (Heer et al., 2017). However, in *moat* mutant embryos, the cell row that starts apical expansion is shifted laterally to row six or seven (Fig. 7E, F, Movie 10). Meanwhile, the middle rows constrict but not as much as those in wild type embryos. This indicates the flanking mesoderm cells ectopically undergo apical constriction, which may stretch the middle cells. We quantified the cell row that transitions from apical constriction to expansion. While the transition point occurs between rows four and five in wild type embryos, it moves laterally to rows five and six in *moat* mutants (Fig. 7G).

Since the gradient of apical constriction is essential for efficient ventral furrow formation (Heer et al., 2017), we examined whether the disrupted gradient of constriction in *moat* mutants could reduce infolding efficiency. Initially, both wild type and *moat* mutant mesoderm cells converge towards the ventral midline as the apical areas of the cells in the middle rows reduce, although the cells in the *moat* mutant embryo constrict more evenly across ventral-lateral axis (Fig 8A, B; Movie 11). After about 5 minutes, flanking mesoderm cells in wild type embryos continue to move towards the midline as middle cells constrict even more, leading to the formation and deepening of the furrow (Fig 8C, upper panels, arrows showing depth; Movie 11). Meanwhile, in *moat* mutant embryos, the cell movement towards the midline stalls and only a shallow furrow forms. It takes much longer for the furrow to reach a similar depth (Fig 8C, lower panels; Movie 11). On average, *moat* mutant mesoderm takes 2.5 min longer than wild type to form a furrow 5 um deep (Fig 8D). Thus, in *moat* mutant mesoderm, the lateral expansion of apical constriction results in uncoordinated and prolonged infolding. Taken together, our data argue the downregulation of junctional Baz in wild type ventral furrow may ensure an optimal gradient of apical constriction and therefore robust internalization of mesoderm.

**Fig. 8.**
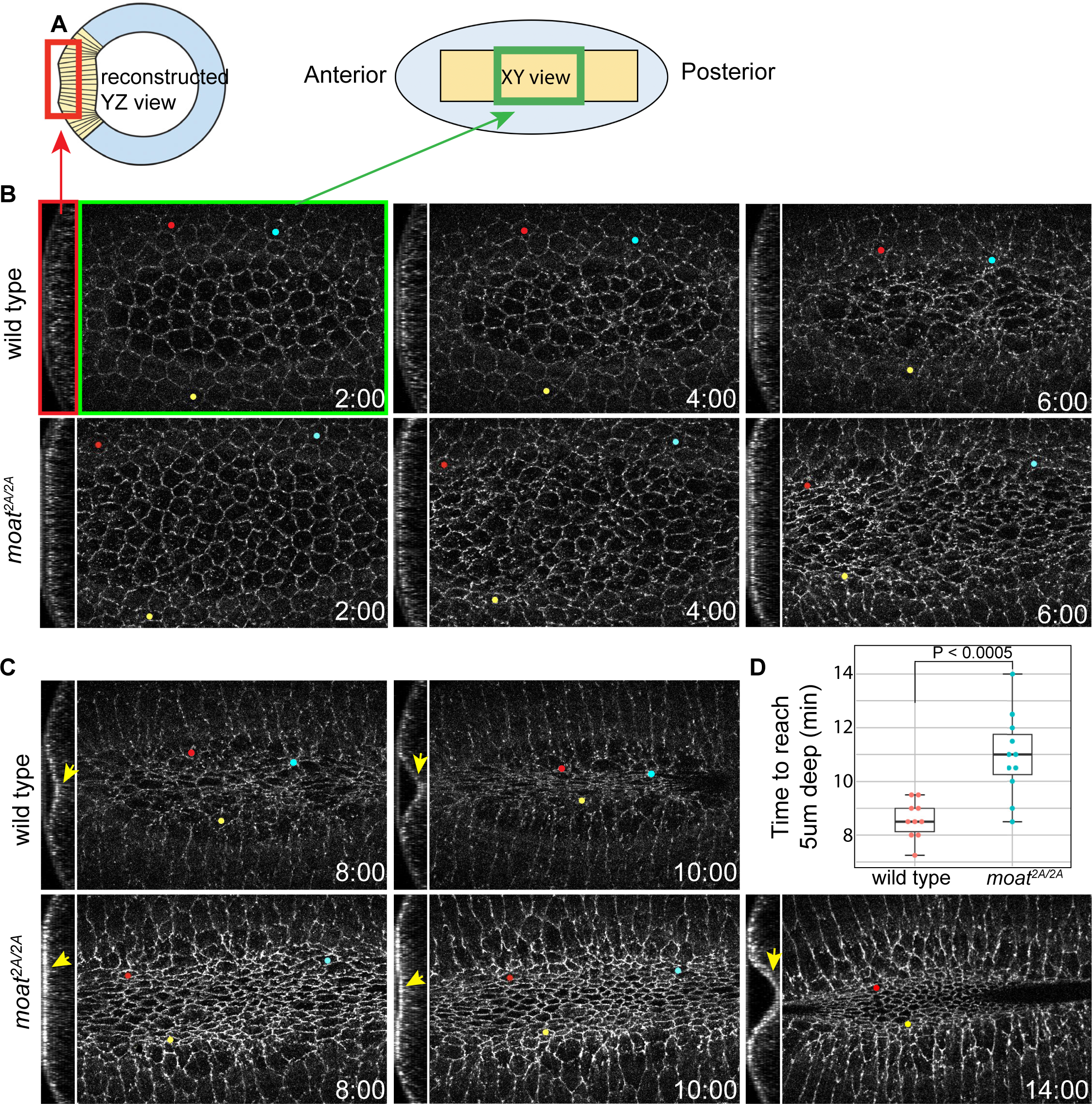
Epithelial folding is prolonged in *moat* mutant mesoderm. (A) Diagram illustrating the two views of constricting mesoderm shown in B and C. Left: cross section view (YZ) showing the depth of ventral furrow. Right: en face view (XY) showing the apical cell outlines. Mesoderm is in yellow. (B-C) Still images from time-lapse movies of folding mesoderm with YZ and XY views as illustrated in A. Tracked cells are marked with colored dots throughout the time. The YZ slices were obtained using the Reslice function in ImageJ. (N) Quantification of the time it takes for the apical surface of the furrow to reache 5um deep. Wild type: N=10; *moat^2A/2A^*: N=11. p=0.00041 determined using student T-tests (two tails, two-sample unequal variance).

## DISCUSSION

The engagement between contractile myosin and adherens junctions executes cell shape changes during many morphogenetic events. However, how the tissue patterning of these two factors determine the tissue boundaries of morphogenetic movement is not well understood. Here, we used the various tissues within the Sna-expressing zone of fly early embryo to investigate morphogenetic boundaries. To identify the boundaries, we clarified the tissue segments within Sna-expressing zone from anterior to posterior: 1) Hkb(+) Gt(-) ectoAMG that participates in neither ventral furrow nor stomodeum invagination; 2) the uncharacterized Hkb(-) Gt(+) cells that forms the anterior end of ventral furrow; 3) endoAMG and 4) mesoderm, both Hkb(-) Gt(-) and forming the rest of the ventral furrow.

With this system, we identified a novel Drosophila gene *moat* that modulates the tissue-specific distribution of adherens junctions through limiting the junctional localization of polarity protein Baz. This Moat-dependent regulation of junction levels is essential for preventing ectopic apical constriction in cells with low-level contractile myosin, thereby maintaining the normal morphogenetic boundaries. We found that in *moat* mutant embryos: along the anterior-posterior axis, the infolding behavior of ventral furrow is expanded to ectoAMG, and along the ventral-lateral axis, the strong apical constriction behavior of middle mesoderm cells is expanded to the flanking mesoderm cells.

Our results propose a model where the combination of different levels of contractile myosin and adherens junctions determines the levels of apical constriction and the subsequent cell shape changes (Fig. 9). We showed that the landscape of adherens junctions is shaped in a two-step mechanism. First, the Baz-dependent adherens junctions uniformly distributed across the embryo are specifically downregulated in the entire Sna-expressing zone (Fig. 1, Fig. 9, checkered green). Next, with the clearance of Baz-dependent adherens junctions, a new distribution of adherens junctions arises in this zone in response to the gradient of contractile myosin: a gradient of adherens junctions forms specifically in ventral furrow cells, whereas junction levels in ectoAMG remain to be low (Fig. 9, green). The gradients of contractile myosin and adherens junctions together lead to a sharpened gradient of apical constriction and robust infolding of ventral furrow. Whereas, the low levels of both contractile myosin and adherens junctions exclude ectoAMG from ventral furrow. In *moat* mutant ectoAMG and flanking ventral furrow cells, the low-level contractile myosin combined with the abnormally high levels of Baz-dependent adherens junctions pushes the level of apical constriction above the threshold for epithelial folding (Fig. 9, blue). We showed that Baz is essential for the ectopic folding in *moat* mutant ectoAMG. This is distinct from normal ventral furrow, where Baz-dependent junctions are less critical since the strong contractile myosin induces high levels of myosin-dependent junctions.

**Fig. 9.**
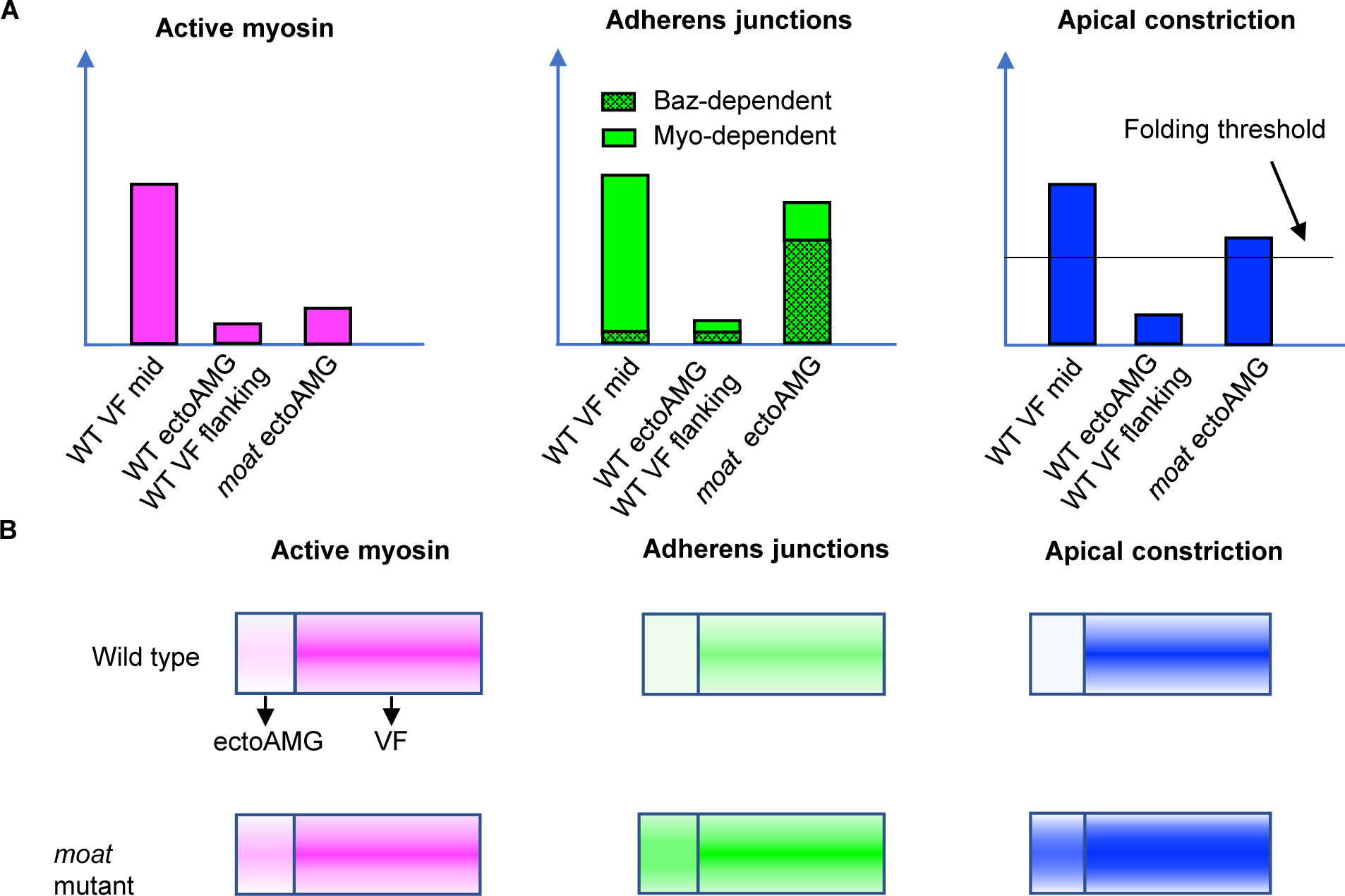
Combination of differential levels of contractile myosin and adherens junctions determines the level of apical constriction. (A) Plots illustrating the relative levels of contractile myosin, adherens junctions and apical constriction in different tissues. (B) Landscapes of the levels of contractile myosin, adherens junctions and apical constriction in Sna-expressing zone.

Moat appears to facilitate Sna in downregulating junctional Baz by reducing the junctional localization of Baz. Past studies of Sna-dependent Baz downregulation raise the question of what is the benefit of weakening junctions before apical constriction, a process that requires strong junctions. Our model suggests the clearance of Baz-dependent adherens junctions in the entire Sna-expressing zone is essential in reestablishing a new landscape of adherens junctions that matches that of contractile myosin. In ectoAMG, this reduces the chance of ectoAMG undergoing ventral furrow-like infolding, especially when the suppression of contractile myosin by Hkb is incomplete. In mesoderm cells, this maximizes the gradient of apical constriction and increases the folding efficiency. Moat could represent the module that optimizes the levels of adherens junctions in the scheme of fast epithelial folding. *moat*, along with genes such as *fog*, *T48,* and *fruhstart* that specifically function in ventral furrow cells at gastrulation stage, ensure successful infolding of mesoderm (Urbansky et al., 2016). They provide us opportunities to understand the shaping of epithelial folds.

What is the molecular and cellular mechanism by which adherens junction levels enhance apical constriction? One possibility is through increasing the effective interaction between actomyosin filaments and cell cortex. Not all contractile actomyosin filaments in the cell are engaged with adherens junctions to constrict apical cortex. An increase in the number of junction puncta and the amount of Cadherin-Catenin complexes per puncta could increase the frequency of actomyosin filament engagement. Another possibility is that high levels of adherens junctions may themselves function as a platform to increase actomyosin assembly or recruitment of myosin activators. Answering this question will shed light on the mechanism of adherens junctions and myosin interactions.

## Acknowledgement

We thank Eric Wieschaus, Chris Doe, Andreas Wodarz and Maria Leptin for fly stocks and reagents. We also thank Eric Wieschaus for supporting the genetic screen. We are grateful for Bloomington *Drosophila* Stock center and BACPAC Resources center for fly strains and vectors, Developmental Studies Hybridoma Bank for antibodies, Flybase for valuable information. We thank Jesus Guerrero for the initial characterization of *moat* mutants and Moat-HA embryos. We thank UNLV Imaging Core for imaging facilities. We thank Laurel Raftery for discussion and comments on the manuscript and Weng lab members for discussion and feedback.

## METHODS

### Fly stocks

Fly lines are listed in table S1

### Generation of fly strains

*moat* mutant allele was generated using CRISPR technique. Two guide RNA fragments were selected: one in the 5’-UTR close to the start codon (gRNA-5) and the other in the exon close to the stop codon (gRNA-3). The phosphorylated gRNA oligos were annealed and cloned into pU6-BbsI-ChiRNA vector (flycrispr.org). For the donor template construct, a 901bp 5’ homology arm and a 1017bp 3’ homology arm were amplified and cloned into a pHD-DsRed vector 5’ and 3’ MCS via enzyme restriction, respectively. The oligonucleotides for gRNA and primers for donor constructs are listed below:

gRNA-5 sequence: 5’-CATGCGGTACGGGTGATTCC**AGG**-3’

Sense oligo: 5’-cttcGCATGCGGTACGGGTGATTCC-3’

Antisense oligo:5’- aaacGGAATCACCCGTACCGCATGC-3’

gRNA-3 sequence: 5’-GCGTGTGCCCGGCTAGCAAT**TGG**-3’

Sense oligo:5’-cttcGCGTGTGCCCGGCTAGCAAT-3’

Antisense oligo:5’-aaacATTGCTAGCCGGGCACACGC-3’

5’ homology arm: F: 5’-TGCACA*gaattc*TGAGATCCCACGTGTTCGTC-3’ (*EcoRI*)

R: 5’-TGCACA*cggccg*GCATGCGGCTACCTGATTAGT-3’ (*EagI*)

3’ homology arm: F: 5’-TGCAGC*actagt*AATTGCTAGCCGGGCACAC-3’ (*Spel*)

R: 5’-TGCAGC*ctcgag*CGCAATCAGTTAATGGGCTGG-3’ (*Xhol*)

The donor and two gRNA constructs were injected into w*; FRT40A; nos-Cas9 attP2 strain by Rainbow Transgenic Flies. The flies carrying the deletion were confirmed by dsRed fluorescence in fly eyes and by PCR using adult fly genomic DNA as a template. The primers for PCR: F: 5’-CACGGCTAGACGGCATTTCT- 3’ and R: 5’-GCAGCTCATGAAAGTCAGGC-3’.

To generate moat rescue flies, the bacterial artificial chromosome (BAC) carrying a small genomic DNA fragment covering *moat* and *nullo* (CH322-67O8) was obtained from the BACPAC Resources Center (BPRC). Embryo injection was done by the BestGene using y^1^w67^c23^; p{CaryP}attP40 strain.

### Embryo fixation and immunostaining

Drosophila embryos were collected on apple juice agar plates at 25°C for 2 hours, followed by an additional 2-hour aging period at the same temperature after the removal of adult flies. Then the embryos were dechorionated with 50% of 8% household bleach (4% sodium hypochlorite) and fixed by heat-methanol protocol as described before (Muller and Wieschaus, 1996). Briefly, in a 15ml glass vial, the dechorionated embryos were incubated in 3ml of hot salt solution (0.4% NaCl, 0.03% Triton x-100) for 10s, and then swiftly cooled by adding 2 volumes of pre-chilled salt solution. Discarded the salt solution from the vial and then added a 1:1 mixture of methanol and heptane. Vortexed vigorously for at least 30 seconds to remove the vitelline membranes. Then, the embryos were transferred into an Eppendorf tube washed with methanol 3 times. The embryos were stored in methanol at -20℃ until use.

Embryos were incubated in 10% BSA blocking buffer for 1 hour, followed by overnight staining with primary antibody at 4℃ and subsequent staining with the secondary antibody for 2 hours at room temperature. The primary and secondary antibodies used in this study are listed in Table S2. Following antibody staining, embryos were sorted and mounted in Aqua-PolyMount (Polysciences). Images are acquired on a Zeiss LSM800 confocal microscope equipped with high-sensitive GaAsp detectors. LD LCI plant-apochromat 25x/0.8 objective was used for whole embryo images and plan-Apochromat 63x/1.4 NA Oil DIC M27 objective was used for high magnification images.

### Live imaging

All images were acquired on the Zeiss LSM800 confocal microscope described above. An LD LCI plant-apochromat 25x/0.8 NA oil objective was used for the anterior midgut imaging and Baz quantification. A Plan-Apochromat 63x/1.4 NA Oil DIC M27 objective was used for E-Cad quantification and mesoderm cell tracking. Diode 488 and 561 nm lasers were used to excite GFP and mCherry respectively. The pinhole is set at 1 Airy unit for 488 for all images. Zeiss Definite Focus was used to maintain the focal plane.

For mesoderm regions, embryos were prepared using the protocol as described (Gu and Weng, 2021). To quantify Baz and E-Cad in the mesoderm (Fig. 1), images were captured at a lower temporal and spatial resolution to minimize photobleaching: a temporal resolution of 1 min per z-stack, a z step of 1 µm and a xy resolution of 0.25 µm and 0.2 µm per pixel, respectively. Baz high magnification images were obtained at a temporal resolution of 30 s per z-stack, a z step of 0.5 µm, and an xy resolution of 0.13 µm per pixel.

For the anterior midgut, embryos were mounted and imaged differently in order to capture most of the ectoAMG on the highly curved anterior end of the embryo with high quality. Embryos were tilted along the anterior-posterior direction during mounting so that the anterior quarter of the embryo landed on the cover glass. Images were acquired with a xy resolution of 0.25 µm per pixel, a z-step of 1 µm, a total depth of 37 µm, and a temporal resolution of 30 s per z-stack.

For cell tracking and apical area measurements, images were captured at a xy resolution of 0.2 µm per pixel, a z-step of 1 µm, a total depth of 18 µm, and a temporal resolution of 30 s per z-stack.

### Scanning EM

Embryos were dechorionated with 50% of 8% household bleach (4% sodium hypochlorite) and fixed for 25 min with 25% glutaraldehyde in 0.1 M sodium cacodylic buffer and heptane. The vitelline membrane was then manually removed in PBS with a needle, and embryos were dehydrated by a gradient of ethanol concentration (25%, 50%, 75%, 95%, and 100%). Embryos were then incubated for 10 min in a 1:1 mixture of ethanol and Hexamethyldisilazane (HMDS), followed by two additional incubations with 100% HMDS. After HMDS evaporated completely, embryos were transferred to the SEM stub and gold coated using a Sputter Coater 108auto (Cressington). Samples were imaged using Hitachi TM-1000.

### Imaging processing and analysis

All images for publication were processed in ImageJ (http://rsb.info.nih.gov/ij/). Brightness and contrast were adjusted for the whole image. Quantitative analysis was done using MATLAB (MathWorks).

To quantify the spot adherens junctions and junctional Baz (Fig. 1), a region of about 9×15 mesoderm cells was used for quantification. Images were thresholded to exclude the uniform and weak membrane or cytoplasmic signals. The threshold is based on the z section 7 µm from the embryo surface, where there is little junctional Baz or spot adherens junctions. The threshold is calculated as pixel value mean plus seven standard deviations for E-Cad::GFP images and mean plus three standard deviations for Baz::GFP images. The difference in the threshold calculation is mainly due to the difference in the distribution of the two proteins outside the puncta pool. While non-puncta E-Cadherin diffuses in the membrane with little cytoplasmic localization, non-puncta Baz is evenly distributed in the cytoplasm with no membrane localization. This leads to much higher non-puncta Baz levels. The mask generated from the initial thresholding is further processed to remove objects (groups of connected pixels) smaller than 4 pixels to generate the final mask for quantification (Fig. S1A). The intensity curve was normalized by the mean intensity of different samples at the starting time point.

For ventral apical area tracking (Fig. 7), live imaging stacks were first flattened to correct the cell tilting caused by embryo curvature using HWada plugin tool in ImageJ (Housei Wada et al,2020). The flattened images were used for cell tracking performed by a MATLAB package, Embryo Development Geometry Explorer (EDGE) (Gelbart, et al., 2012). EDGE detects the xy coordinates of cell vertices which are used for segmenting and tracking cells. Cell segmentation errors were manually corrected. The section at 4 µm deep from the apical surface was used to calculate apical area because the severe deformation of cell outline in the more apical sections prevents accurate segmentation and tracking. To group cells in rows, we defined the middle two cell rows using the time points when the furrow was a few microns deep and traced the cell rows back to early time points. We plotted the average apical area of all cells in each row over time. The apical areas of individual rows are normalized by the areas of corresponding row at the starting time point to show the fold change.

The intensity of spot adherens junctions in Fig. 7 was measured in a similar way as that in Fig. 1. The cell rows were defined from the cell tracking. To quantify E-Cadherin accurately, embryos that were mounted slightly off-center relative to the ventral midline were used for quantification and only 6 rows on one side of the embryo were plotted. This is because the fluorescence from the cells at the position that directly touch the coverslip is less refracted and artificially stronger than that from the cells positioned on the flanking slopes of the embryo. The gap between the slope of the embryo and the coverslip leads to loss of signals.

### Quantification of embryo phenotype

The fixed embryos, stained with Nrt or Arm, or imaged by SEM, were manually categorized based on their phenotype.

For scoring ectopic ectoAMG furrow phenotype, embryos of late stage 6 and stage 7 were used. Embryos with putative ectoAMG cells exhibiting relaxed apical surfaces that follow the curvature of the embryos were classified as having a normal end. Embryos with putative ectoAMG exhibiting apical constriction and forming a furrow were defined as having an extended end. (Fig. 2B, Fig. 6C) For scoring ventral furrow closure phenotypes (Fig. 6D), embryos were sorted by whether the middle ⅓ of the ventral furrow is closed. Embryos from late stage 6 and stage 7 were used for quantification. Ventral furrows in wild type embryos usually close by late stage 6 and become less distinguishable by stage 8 when domain 14 enters cell cycle.

### Western blot

Drosophila embryos were collected on apple juice agar plates at 25°C for 2 hours, followed by an additional 2-hour aging period at the same temperature after the removal of adult flies. Then the embryos were dechorionated by 50% of 8% household bleach (4% sodium hypochlorite). Embryo extraction was conducted by incubating dechorionated embryos in 2x Laemmli buffer (Bio-Rad) with 5% 2-mercaptoethanol at 95°C for 2 min. The lysates were then cleared by centrifugation at 16,000g for 5 min. 8% surePAGE gels (GenScript) were used for electrophoresis and Immobilon-FL PVDF membranes (Millipore) were used for transfer. Membranes were blocked for 1 h in PBST (0.1% Tween-20 in PBS) containing 10% dry milk. Then the membranes were incubated in primary antibody overnight at 4°C (Guinea pig anti-Baz, 1:500; Mouse anti-Tubulin, 1:2000) and subsequently in secondary antibody for 1 hour at room temperature (IRDye-coupled secondary antibody: Donkey anti-Gp 800CW Cat# 926-32411 and Goat anti-Mouse 680RD Cat# 926-68070, 1:5000). Membranes were visualized with an Odyssey infrared imaging system (LI-COR Bioscience), and brightness and contrast were adjusted for the whole image using ImageStudio software.

### RNA in situ hybridization

The probe template was generated using the following primers: Forward: 5’- TGCAGAACAGTCGACAAGGA-3’ and Reverse: 5’-

GCTGGGCAATGCAATCGTAGC-3’. DIG-labeled antisense and sense probes were synthesized using T7 and SP6 RNA polymerase following the instructions provided in the MGEA script kit from Applied Bio. For hybridization, the eggs were dechorionated in 4% sodium hypochlorite for 1 min, fixed for 20 min in a 1:1 mixture of heptane and 4% paraformaldehyde in PBS, and devitellinized in 1:1 methanol/ heptane mixture and washed in PBST. In situ hybridizations on whole-mount fixed eggs were essentially performed as described before (Tautz, D. & Pfeifle, C, 1989). Then the embryos were blocked and incubated with primary antibody (anti-DIG-AP, Millipore Sigma Cat#11093274910) for 2 hours at RT. For color development reaction, the embryos were incubated in NBT/BCIP solution (Millipore Sigma, Cat#11681451001) at RT in the dark. The reaction was stopped by washing several times in PBS with 0.1% Tween 20. Embryos were photographed using a Nikon 90i Eclipse microscope with S Fluor 10x DIC N1 optics and a motorized stage.

### Statistical analysis

Statistical analyses in Fig. 7 and 8 were conducted using a two-tailed unequal Student’s t-test.

